# A complete mitochondrial genome of a Roman-era *Plasmodium falciparum*

**DOI:** 10.1101/2024.03.05.583465

**Authors:** Alejandro Llanos-Lizcano, Michelle Hämmerle, Alessandra Sperduti, Susanna Sawyer, Brina Zagorc, Kadir Toykan Özdoğan, Meriam Guellil, Olivia Cheronet, Martin Kuhlwilm, Ron Pinhasi, Pere Gelabert

## Abstract

Malaria has historically been one of the leading infection-related causes of death in human populations. To this day, it continues to pose a significant public health threat in African countries, particularly among children. Humans are affected by five *Plasmodium* species, with *Plasmodium falciparum* being the most lethal. The study of pathogenic DNA from ancient human remains has been vital in understanding the origin, evolution, and virulence of human-infecting pathogens. However, there have been no complete pre-20th century mitochondrial DNA (mtDNA) or genomic sequences of *Plasmodium falciparum* reported to date. This gap in knowledge makes it difficult to understand the genetic dynamics of this pathogen in the past. The difficulty in identifying ancient malaria cases through bioarchaeology and the infrequent presence of *Plasmodium* DNA in ancient bones contribute to these limitations. Here, we present the first complete mtDNA genome of *P. falciparum* recovered from an archaeological skeleton (a 2^nd^ century CE Roman individual from Italy). The study of the 43-fold mtDNA genome supports the hypothesis of an Indian origin for *P. falciparum* in Europe and provides evidence for the genetic continuity of this lineage over the past 2,000 years. Additionally, our research highlights that extensive sampling may be necessary for malaria screening to gain insights into the evolution of this vector-borne disease from archaeological samples.

## Main

Malaria is an infectious disease caused by various *Plasmodium* species. Among those affecting humans, *Plasmodium vivax* has the broadest distribution, while *Plasmodium falciparum* is responsible for the majority of malaria-associated deaths. Currently, malaria’s geographic distribution predominantly spans warmer climates near the equator, covering Africa, parts of the Middle East, Southeast Asia, China, and the Americas. Due to its extensive reach and severity, malaria remains one of the most significant health threats to humans. It is estimated that in 2016, approximately 455,000 people died from malaria, with 91% of these deaths occurring in Africa ^1^. Both *P. vivax* and *P. falciparum* are believed to have originated in Africa around 50,000-60,000 years ago^2,3^ and spread worldwide with complex patterns of migration, probably following human migrations^2,4^. Into the 20^th^ century, malaria was widespread worldwide with its widest known geographical distribution, including southern and northern Europe and the USA^5,6^.

It has been hypothesised that both *P. vivax* and *P. falciparum* may have reached Europe during the Neolithic, about 8,500 years ago, due to a combination of favourable parameters, including climatic conditions, increased human population densities, and the presence of a capable *Anopheles* vector species^7^. However, this claim lacks archaeological or genetic evidence and remains contested^3^. Nevertheless, there is consensus that *Plasmodium* species have been present in Europe since at least the Roman Imperial period, particularly around the Mediterranean shores of the Roman Empire, coinciding with endemic areas of *Anopheles spp—*endemic areas^7^. Scholars addressing the effects of this endemic disease on societies in antiquity have stressed the dramatic political and economic consequences of malaria, an issue sometimes neglected by historiography^8^. Malaria remained endemic in Europe until the 1970s, extending from the Baltic Sea to the Mediterranean^9^. The lack of ancient malarial genomes, however, leaves the initial spread of malaria to Europe and its origin unclear.

The identification and sequencing of ancient malaria strains present several challenges. Firstly, osteological lesions are not indicative of infection, and archival records confirming infection are often missing from archaeological skeletal collections. As a result, the detection of malaria, similar to HBV and other pathogens, relies on indiscriminate sampling of individuals. Secondly, the success in recovering ancient *Plasmodium* genomes from human remains is contingent on the survival of *Plasmodium* DNA in bone and dental tissues. Some studies have suggested the presence of *P. falciparum* DNA in ancient individuals, such as a 5^th^ century CE infant from Lugnano in Teverina, Italy^10^, and in ancient Egyptian mummies^11^. However, these studies fell short of providing definitive evidence of the pathogens, a gap that next-generation sequencing techniques can now address. A recent study comparing the reliability of antigen detection and DNA sequencing in identifying pathogens in ancient skeletal remains found that, despite a limited sample size, paleogenomics methods are the most dependable for this purpose^12–14^. Microscopy remains an alternative for mummified tissues^15^.

Currently, only two sequences from ancient *Plasmodium* strains have been identified. The first is a complete mitochondrial genome from a mid-20^th^ century CE individual from Spain (Ebro-1944), closely related to contemporary Indian strains^16–18^. This individual had a co-infection of *P. vivax* and *P. falciparum*, a common occurrence in regions where both species are endemic^19^. The genetic similarity between the Spanish sample and Indian strains lends support to the hypothesis of a spread of *P. falciparum* from Asia to Europe, possibly during ancient times. The second sequence is a low-coverage partial genome obtained from two individuals in Italy, LV13 (Velia) and LG20 (Vagnari), dated to the 1^st^–2^nd^ century CE. These sequences are the oldest evidence of *P. falciparum* presence to date and the only pre-20th Century genetic data available^20^. The combined data from these individuals covers 50.8% of the 5,967 bp *P. falciparum* mitochondrial genome. However, due to the low coverage, it is challenging to differentiate between DNA damage, sequencing errors, and actual single nucleotide polymorphisms (SNPs) in these genomes. Thus, while these partial sequences provide initial evidence for the phylogenetic positioning of ancient European *P. falciparum* strains within a modern clade, they lack the comprehensive coverage necessary for high-resolution phylogenetic analysis^20^.

Here, we introduce the first complete mitochondrial genome sequence (43-fold coverage) of *Plasmodium falciparum* from the Roman era, derived from an individual known as Velia-186 (LV13), previously confirmed to be infected with the pathogen^20^.

In an initial screening, we produced sequencing libraries from both teeth and parts of the femur of individual Velia-186 (Velia, Porta Marina necropolis, I-II cent. CE) .We targeted teeth based on evidence that other pathogens are well-preserved in dental tissues^21^ and the femoral diaphysis and head, given that *P. falciparum* gametocytes commonly mature in human bone marrow^22^. As one of the libraries yielded just over 100 unique reads for P. falciparum, we decided to increase the efforts for recovering more DNA. For that purpose, 38 DNA libraries from seven teeth were generated by sampling at least two roots per tooth (see Supp. Table S1). The libraries were enriched with in-solution baits covering both *P. falciparum* and *P. vivax* mitochondrial genomes and sequenced on an Illumina NextSeq 550, generating paired-end reads with a length of 150 bp. After preprocessing, quality control, and collapsing, 946 million, averaging 22 million reads per library (standard deviation (SD) = 1.167 million), were recovered, and the data was subsequently merged. To assess whether the recovered reads originated from a co-infection of *P. falciparum* and *P. vivax*, we performed competitive mapping, which maps reads to different reference genomes simultaneously to ascertain to which genome the read fits best. Competitive mappings for *P. falciparum* and *P. vivax* resulted in 33.66-fold (Fig. 1C, Table S1) and 1.75-fold (Fig. 1C, Table S1) depth of coverage, respectively, indicating that the vast majority of reads stem from *P. falciparum* and not *P. vivax*. We then performed comparative mapping, which maps to each genome individually to obtain the coverage per species. This mapping resulted in a mean depth of 21.61 for *P. vivax*. However, only 22% of the genome has coverage due to uneven mapping, which indicates mismapping. On the other hand, the mapping to P. *falciparum* shows an even distribution of coverage with an average depth of 43.67-fold. (Fig. 1C, Table S1). Based on these results, we deduce that the Velia-186 individual was only infected with *P. falciparum* and exclude the possibility of a co-infection with *P. vivax* as in Ebro-1944. Hence, below, we focus on reads mapped to the *P. falciparum* mtDNA genome. Additionally, we inspected misincorporation patterns characteristic of ancient DNA, finding a clear deamination pattern in both 5’ (22%) and 3’ (18%) ends, corroborating an ancient origin of the sequencing data (Fig. 1C). Furthermore, the read-length distribution shows the typical small fragment lengths associated with ancient DNA (Supp. Table S1).

**Figure 1:**
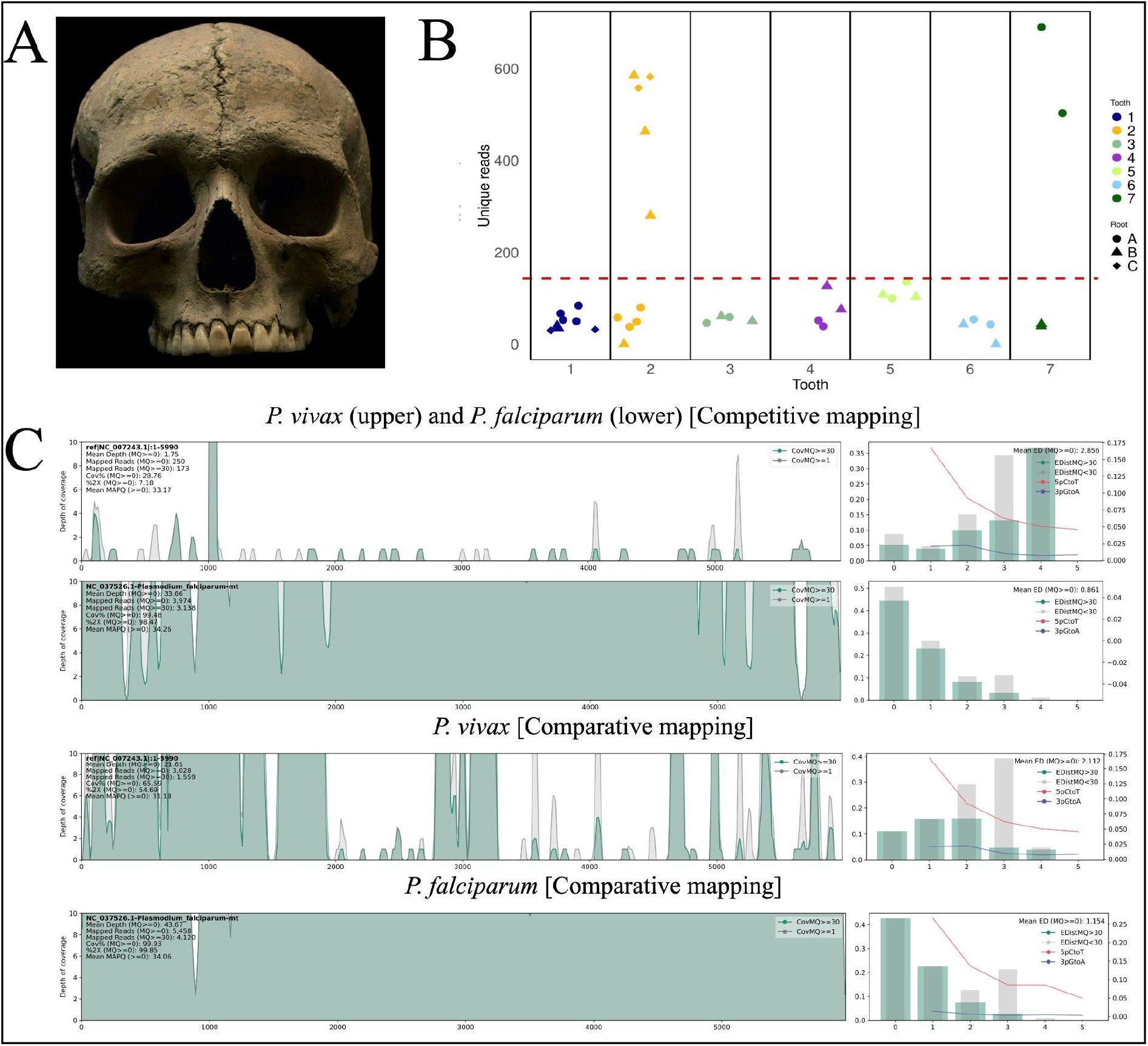
(A) Velia-186 skull (male 20-25 years old). Photo by the Museum of Civilizations. (B) The number of unique reads mapping to the reference genome K1 [NC_037526] was recovered from each sequencing library. The red line denotes the average amount of reads recovered per library. Samples 2, 3 (Upper right second molar), and 8 (Lower right first molar) contribute the majority of reads. (C) Mapping plots to both P. vivax and P. falciparum. Reads with an MQ of or above 30 are depicted in green. The coverage is shown across the whole mitochondrial genome. The bar plot on the right depicts the edit distance and the percentage of C -> T mutations at the 5`end and the G -> A mutations at the 3’end.

In our study, following the identification of a solitary *P. falciparum* infection, we proceeded to analyze the *P. falciparum* DNA across various sequencing libraries. This analysis was centred on aligned reads with mapping scores above 30 (refer to Supp. Table S1). Notably, we found no reads aligning to *P. falciparum* on the external surfaces of the first right inferior premolar nor in the femoral diaphysis or head (Supp. Table S1). On average, each of the libraries from the seven teeth produced 108 reads (with a standard deviation of 180 across all 38 libraries). Intriguingly, a mere seven out of the 38 libraries contributed to 72% of the unique fragments (see Fig. 1B).

To explore any potential correlation between the dental samples and their respective sequencing yields, we conducted an analysis but found no significant correlation (R^2^ = 0.07 and p = 0.65). Further, considering the non-normal distribution of reads, we investigated whether there was greater diversity within the teeth samples compared to between them. This was assessed using a Kruskal-Wallis test (χ2(6) = 11.06, p = 0.08), which revealed no significant differences in medians, suggesting homogeneity across the dental samples.

These findings underscore a considerable intra-individual variation in the presence of *Plasmodium* within a single tooth. This variation implies that for effective pathogen sampling, a strategy encompassing multiple samples from an individual might be more optimal than relying on a single sample.

We combined the 5,458 mapped reads from all seven teeth, which gives us a mitochondrial genome of *P. falciparum* with an average depth of 43-fold. We used these reads to generate a consensus sequence with ANGSD (version 0.941) and obtained a mitochondrial genome with 99.1% of the sequence covered. The consensus sequence has three gaps with a size of 59 (858-917), 20 (1151-1171) and 9 (3495-3504) base pairs. In a previous study on the same individual, based on only a few sequenced fragments and low depth of coverage, Marciniak et al. 2016 reported 21 mutations between the ancient genome and the mitochondrial *P. falciparum* reference genome. The SNPs in our consensus sequence were inspected individually in IGV (v2_16.0) to verify and validate their presence. Our high-coverage genome consensus sequence has seven mutations compared to the reference strain mitochondrial genome. When comparing our high-coverage calling with the previously reported mutations in LV-13^20^ we could only verify two of the previously reported mutations (2763, C > T; 3938, A > T) while adding five well-supported ones (74, A > T; 276 G > A; 725, C > T; 772, T > C; 2172, T > C), all supported by at least ten reads (see Supp. Table S2). The previously reported higher number of mutations is probably a result of the low coverage of degraded DNA. Four of the seven mutations are located in coding regions of the mtDNA (Supp. Table S2). However, none of these is non-synonymous using the Apollo^23^ annotation tool accessed through PlasmoDB^24^. More details about the mutations are presented in Supplementary Table S2.

To elucidate how our *P. falciparum* consensus sequence compares to other *P. falciparum* genomes, we downloaded 339 mitochondrial genomes of *P. falciparum* strains from NCBI, representing the entire present-day diversity of *P. falciparum*. (see Supp. Table S3). First, we observed that only two substitutions, at positions 2172 (T > C) and 3938 (A > T), were not present in any *P. falciparum* strain of the dataset. Next, we performed a maximum likelihood tree using a multiple sequence alignment, including our ancient consensus sequence (n = 340). Although some clades cluster geographically, we observed that the current mtDNA genetic diversity does not exclusively reflect a geographical distribution, as previously reported ^25^. In the phylogenetic tree, Velia-186 clusters exclusively with strains currently found in India with 95 bootstrap support. (Fig. 2, Supp. Fig. 1). Out of the seven SNPs described above, mutations 276 and 2763 are observed in the present-day Indian strains from different locations in India^21^, which supports the clustering of the Velia-186 sequence close to the Indian strains. The SNP at position 725 (C>T), combined with the 276 and 2763 mutations, is characteristic of the Indian subclade called PfIndia, described by Tyagi et al., 2014 ^26^. These three defining mutations also exist in Ebro-1944, indicating genetic similarity between European

**Figure 2:**
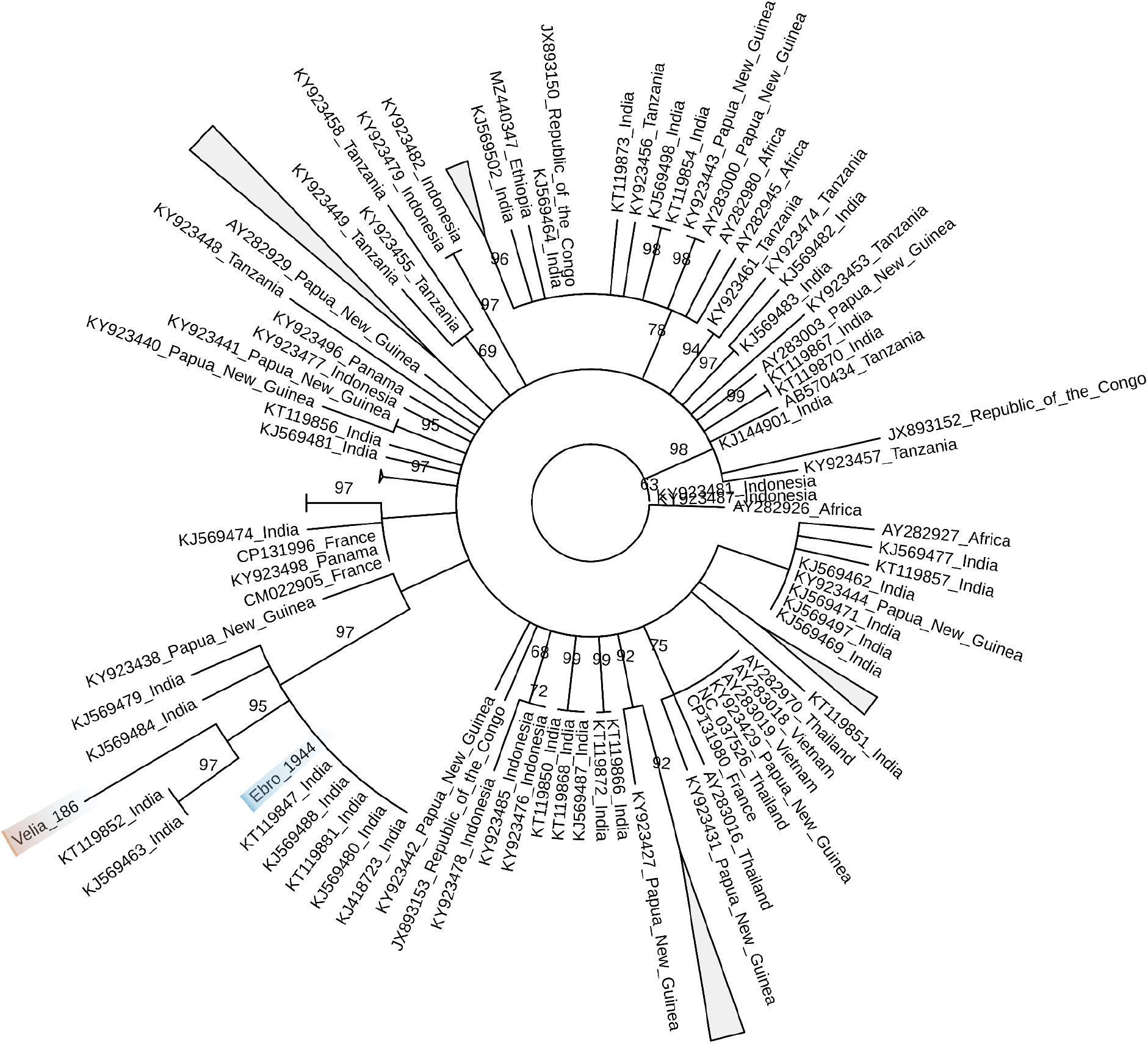
*P. falciparum* mtDNA maximum likelihood phylogeny. It is observable that the *P. falciparum* from Velia-186 in red clusters with Indian strains and clusters close to Ebro-1944 in blue. The tree was visualised with TreeViewer ^28^. We present an unrooted phylogeny without branch lengths. Numbers represent bootstrap values.

samples. As this combination of SNPs is also evident in *P. falciparum*-like pathogens infecting monkeys, it is suggested to be the missing link between animal- and human-infecting parasites^26,27^.

Our findings lend further credence to the hypothesis proposed by de-Dios et al., 2019, regarding the Asian origin of *P. falciparum* in Europe during ancient times. This hypothesis suggests that the parasite might have disseminated across Europe during the Achaemenid Empire and the subsequent Hellenistic period, propelled by the extensive movement of goods and people from various Asian regions to Europe. Remarkably, despite the Velia-186 mitochondrial genome originating from an individual nearly two millennia old, its sequence demonstrates a close phylogenetic relationship with the Ebro-1944 *P. falciparum* mitochondrial genome from 1940s Spain. This connection implies a genetic continuity of the parasite in Europe over the last 2000 years. Additionally, the partial nuclear data from the Ebro-1944 sample indicated an affinity with Asian strains, as previously suggested by the mtDNA^16,18^. However, since nuclear data from Velia-186 has not yet been obtained, it raises the intriguing question of whether the nuclear genome of the Velia-186 strain would similarly exhibit an affinity with Asian strains.

## Author Contributions

P. G. conceptualised the study. A. S. sampled and provided archeo-anthropological context, A. L-L, O. C and S. S. performed the experiments, P. G., M. G., A. L-L., and M. H. analysed the data, and P. G., A. L-L., R. P, A. S, M. K, M. G, S.S. and M. H. wrote the text with inputs from all collaborators.

## Acknowledgements

The study was funded through FWF Principal Investigator grant P–36433 and INFRAVEC ISID_2019 to P. G. Alejandro Llanos-Lizcano was funded by Colfuturo (Fundación para el futuro de Colombia) during his MSc. This project has been funded by the Vienna Science and Technology Fund (WWTF) [10.47379/VRG20001] and by the Austrian Science Fund (FWF) [FW547002] to M.K.

## STAR Methods

### Key resources table

| REAGENT or RESOURCE | SOURCE |
| --- | --- |
| **Biological Samples** |  |
| Velia_gr. 186 Lower premolar left 1 | Museo della Civiltà |
| Velia_gr. 186 Upper molar right 2 | Museo della Civiltà |
| Velia_gr. 186 Upper molar left 1 | Museo della Civiltà |
| Velia_gr. 186 Lower molar left 2 | Museo della Civiltà |
| Velia_gr. 186 Upper molar right 1 | Museo della Civiltà |
| Velia_gr. 186 Lower molar left 3 | Museo della Civiltà |
| Velia_gr. 186 Lower molar right 1 | Museo della Civiltà |
| Velia_gr. 186 Head of the femur | Museo della Civiltà |
| Velia_gr. 186 Distal epiphysis of Femur (Left) | Museo della Civiltà |
| Velia_gr. 186 Lower premolar (Left) (Cavity) | Museo della Civiltà |
| Velia_gr. 186 Surface of the root of the lower premolar | Museo della Civiltà |
| Deposited Data |  |
| One mitochondrial whole genome from a Roman individual | This paper |
| SOFTWARE AND ALGORITHMS |  |
| Cutadapt 4.5 | 29 |
| PEAR 0.9.11 | 30 |
| BWA 0.7.17 | 31 |
| Samtools 1.9 | 31 |
| Picard Tools 3.1.1 | 32 |
| aDNA-BAMplotter | 33 |
| ANGSD 0.941 | 34 |
| IGV 2.16.1 | 35 |
| MAFFT v7.520 | 36 |
| IQTree2 2.2.6 | 37 |
| Haplogrep 3.2.2.1 | 38 |

### Lead contact

Questions regarding further information on materials, datasets, and protocols should be directed to and will be fulfilled by the Lead Contact, Pere Gelabert.

### Materials availability

The raw genomic data used in all the analyses can be accessed at the European Nucleotide Archive (ENA) under the accession number: PRJEB72667.

## Data and code availability

Sequencing data and the filtered sequences are available at the European Nucleotide Archive (ENA) under the accession number: PRJEB72667. All code used in this study and other previously published genomic data is available at the sources referenced in the key resource table.

## Method Details

### Archeological context

We sampled seven dental pieces from Individual Velia-186. Velia is located on a peninsula on the Tyrrhenian coast 112 km southeast of Naples in the region of Campania and was an important port during Roman times. Several bioarchaeological analyses have been carried out on individuals from this cemetery ^41–43^, and *P. falciparum* ^*20*^ was identified in the individual we selected for downstream analyses. This sample is from the 1^st^-2^nd^ centuries CE.

Velia is located on a peninsula on the Tyrrhenian coast 112 km southeast of Naples and was incorporated into the Roman territory in the III century. BCE became a port city utilised for the shipment of goods, boat maintenance, fish processing, and arboriculture. The subsistence was also provided by cultivations in the hinterland and well-watered intramural areas. The cemetery of Porta Marina (I-II cent. CE) was investigated by Fiammenghi (2003) and led to the identification of approximately 330 burials (mostly inhumations). The human skeletal material is entrusted for anthropological study to the Bioarchaeology Service of the Museum of Civilizations based in Rome to reconstruct the funerary rituals and describe the demographic and bio-social characteristics of the ancient inhabitants of Velia within an interpretative framework guided by historical-archaeological evidence. The analyses, still in progress, have involved various aspects, such as sex and age composition of sample ^43^, health status ^44,45^, and prevalent working activities. ^42,46^, diet ^47^, migration and individual mobility patterns ^39,40^, and dental anomalies ^48^

### Experimental model and subject details

Velia-186 is represented by a complete and well-preserved skeleton, lacking only small portions of the skull, part of the scapular blades, and some hand and foot elements. The right iliac fossa presents a large green stain from a bronze object that came into contact with the bone after the soft tissue decomposition. The morphological and morphometric analyses led to a diagnosis of a young adult male (20-25 years). The values of oxygen and strontium isotopes indicate that the individual was likely born and raised in Velia ^39,40^. The skeleton shows a robust morphology with mildly developed muscle attachments. The estimated living height (by Pearson regression formulas on the femur maximum length) is 166.9 cm, slightly above the average of the VELIA male series (164.7±4.8 cm). The distal segment of the diaphysis of the right femur shows evidence of a healed trauma. The direction of the main axis of the shaft is unaltered, but its medial surface presents a slight deformation and thickening corresponding to the callus formation. Velia-186 shows developmental anomalies: retention of the metopic suture; unfused scapular acromion (right side;); bipartite rib (left side); bipartite right patella. The individual presents cribra orbitalia on both orbital roofs. The lesions (Type 3, according to ^13^ ) are in a remodelling phase.

The teeth have moderate wear, with small patches of dentine exposure on the anterior dentition and on the first molars (Stage 3-4) ^14^. The first mandibular left molar was lost antemortem, followed by alveolar bone remodelling and complete filling of the roots’ sockets. The adjacent second molar is affected by a small caries lesion on the mesial aspect of the crown. Very slight supragingival calculus deposits (grade 1 by Brothwell scale) are present on a few teeth, affecting mainly the interproximal surfaces of the lower anterior dentition. The following dental morphological variants were recorded: shovel shape (grade 2) on the upper incisors; dental tuberculum of grade 3 on both the left incisors and grade 4 on the right ones; Carabelli’s cup on both first molars (grade 3); 6th cusp on the third lower molar (grade 3).

### Laboratory procedures

100 mg was ground from multiple locations of different teeth and bone material (Table S1). DNA was extracted from bone powder in the Ancient DNA (aDNA) Laboratory at the University of Vienna following detailed protocols adapted to aDNA ^49^. Single-stranded libraries were prepared following detailed protocols for ancient DNA^50^. Libraries were enriched with an in-solution capture designed by Daiiecel Arbor Biosciences (https://arborbiosci.com/products/targeted-ngs/mybaits-custom-kits/mybaits-custom-dna-seq/)following the manufacturer manual. The solution kit included baits targeting the mtDNA sequences of both *P. falciparum* and *P. vivax*. Libraries were pooled in 20 *μl* and sequenced 2×150 PE on NextSeq 550 at the Polo d’Innovazione di Genomica Genetica e Biologia (Siena, Italy).

### Bioinformatics

Sequenced reads were clipped with cutadapt 4.5 ^29^, removing Illumina adapters and base quality 30, and later collapsed using PEAR version 0.9.11 ^30^, requiring a minimum overlap of 11 bp and a minimum length of 30 bp. Filtered collapsed reads were aligned with a competitive mapping of both *P. vivax* and *P. falciparum* (Salvador 1 [ID: NC_007243.1] and K1[ID: NC_037526], respectively) using BWA 0.7.17 aln ^31^, disabling seeding to allow a higher sensitivity of aDNA reads ^51^, gap open penalty of 2 and edit distance of 0.04. Next, duplicates and low mapping-quality reads were removed using Samtools 1.9 ^52^ and Picard-tools 3.1.1 ^32^. We calculated the deamination pattern using MapDamage ^38^; mapping plots were created using aDNA-BAMplotter ^33^. We used this competitive mapping between *P. falciparum and P. vivax* to determine the possible presence of co-infection.

A consensus sequence was obtained using ANGSD (v0.941) ^53^ using the majority of calls and excluding sites with coverage lower than 5, all the SNPs are fixed. The detected mutations were examined visually with IGV ^35^. The consensus sequence was aligned with multiple *P. falciparum* sequences^16,26,54–59^ with MAFFT (v7.520) ^36^. For the phylogeny, the alignment was filtered for positions with 90% coverage, which left 5884 sites, including variable 100 sites. The TVMe+ASC substitution model was chosen with IQTree2 (v 2.2.6), and we constructed a Maximum likelihood tree with 1000 nonparametric bootstrap replicates.

### Human mtDNA analyses

We aligned the reads against the human genome (hg19), including the mtDNA sequence (rCRS). We could recover 2,141,320 reads by combining all the libraries, which all show consistent signals of deamination due to the age. 153022 out of these reads are aligning to the human rCRS genome. We recover a 700X mtDNA sequence. We recovered the consensus sequence using ANGSD ^53^, and identified the haplogroup using Haplogrep 3.2.2.1 ^60^. We identified that the individual Velia 186 belongs to haplogroup T2b7a (88%), which is already identified in MBA individuals from the Levant ^61^ and is also present in several present-day Mediterranean populations and is mostly present in European populations ^62^.

## Supplementary table titles

**Table S1:** Sequencing results of the samples included in the analysis

**Table S2:** List of mutations in the Velia-186 sequence

**Table S3:** *P. falciparum* mtDNA sequences used in the analyses

## Supplementary Figure

**Supplementary Figure 1:**
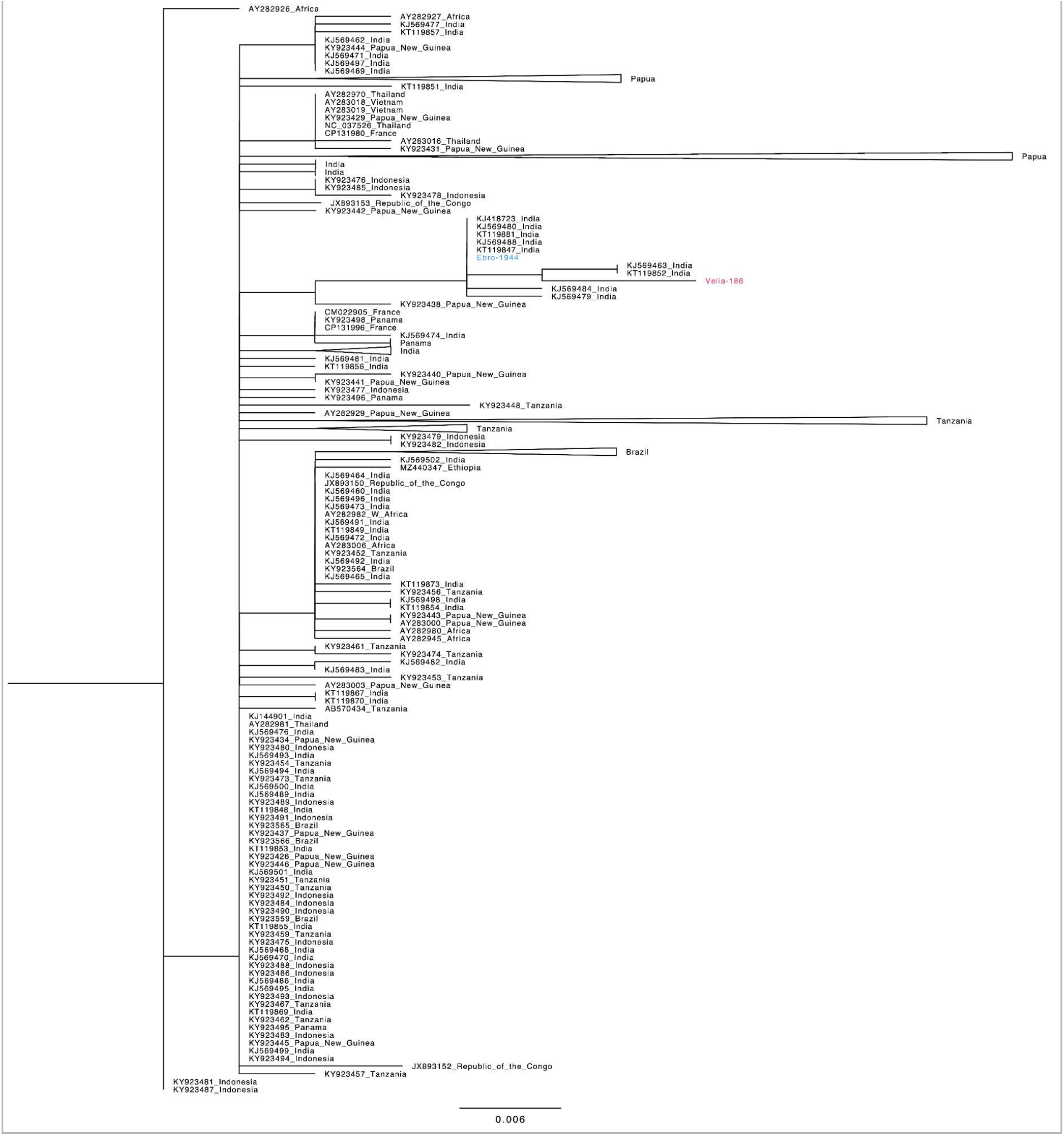
*P. falciparum* mtDNA Maximum Likelihood phylogeny. The sample sequenced in this study is written in red (Velia-186) and the other partial European genome (Ebro-1944) in blue.

